# Predicting Targeted Cancer Therapeutics

**DOI:** 10.1101/057901

**Authors:** Aaron Wise, Murat Can Cobanoglu

## Abstract

**Motivation:** Cancer is a complex and evolving disease, making it difficult to discover effective treatments. Traditional drug discovery relies on high-throughput screening on reductionist models in order to enable the testing of 10^5^ or 10^6^ compounds. These assays lack the complexity of the human disease. Functional assays overcome this limitation by testing drugs on human tumors, however they can only test few drugs, and remain restricted to diagnostic use. An algorithm that identifies hits with fewer experiments could enable the use of functional assays for *de novo* drug discovery.

**Results:** We developed a novel approach that we termed ‘algorithmic ideation’ (AI) to select experiments, and demonstrated that this approach discovers hits 10^4^ times more effectively than brute-force screening. The algorithm trains on known drug-target-disease associations assembled as a tensor, built from the (public) TCGA and STITCH databases and predicts novel associations. We evaluated our tensor completion approach using a temporal cutoff with data prior to 2012 used as training data, and data from 2012 to 2015 used as testing data. Our approach achieved 10^4^-fold more efficient hit discovery than the traditional brute-force high-throughput screening. We further tested the method in a sparse, low data regime by removing up to 90% of the training data, and demonstrated the robustness of the approach. Finally we test predictive performance on drugs with no previously known interactions, and the algorithm demonstrates 10^3^-fold improvement in this challenging problem. Thus algorithmic ideation can potentially enable targeted antineoplastic discovery on functional assays.

**Availability:** Freely accessible at https://bitbucket.org/aiinc/drugx.

## 1 Introduction

Cancer is a disease characterized by genomic instability [10] which gives rise to disease evolution [12] and heterogeneity [20, 23]. These characteristics make cancer therapeutics particularly difficult to discover [25] when drug discovery is already a process with high attrition rates [6, 13, 29]. Commonly used cell line or single target based reductionist approaches cannot accurately account for the tumor’s microenvironment or subclonal/phenotypic heterogeneity [30] thus necessitating more representative assays.

Testing of antineoplastic therapeutic candidates on human tumor tissue alleviates these concerns by generating results that are highly relevant to the human disease context [4, 14, 16, 22, 32, 15]. However there is a key obstacle to the use of human tumor based testing for driving *de novo* drug discovery: the divergence between the numbers of compounds tested in traditional drug discovery campaigns - high throughput screening (HTS) of 10^5^ to 10^6^ compounds [21] - and the number of compounds that can be tested on a human tumor sample - 10^1^ to 10^2^ [22]. If human tumor samples are to be used for screening an entire chemical library, one must either use multiple samples from many individuals or use immortalized cell lines originally derived from primary tumors. Using samples from multiple patients in the same screening would present high batch-to-batch variability problems. When using patient derived cell lines the 3D tumor microenvironment and the clonal heterogeneity are lost while the selective pressure of the culture environment can also deviate the phenotypic responses. Using even more reductionist models such as isolated protein binding assays or enzymatic assays completely lack the intracellular context, in addition to the intercellular context, including some key features such as protein-protein interactions or compensatory signaling networks. Thus no drug discovery pipeline that relies on screening large numbers of compounds can feasibly be run on human tumor samples.

Consequently functional assays that allow for testing drug activity on human tumor samples have so far been restricted primarily to diagnostic use - i.e., determining which of the approved therapy alternatives would work best for a specific patient [7]. However an algorithmic approach that could effectively discover hits with few numbers of tested compounds could utilize functional assays for *de novo* drug discovery. We have addressed this important challenge and hereby present our methodology for computationally predicting targeted cancer therapeutics. Our approach, which we termed ‘algorithmic ideation’, relies on the factorization of a tensor representation of drug-target-disease associations.

New machine learning advances in analyzing the relationships in large networks for predictive purposes such as probabilistic matrix factorization [24], restricted Boltzmann machines [28], probabilistic tensor factorization [35], and factorization machines [26, 27] have provided a rich algorithmic basis for large-scale analysis of genomics and drug-discovery questions. Some recent work applying matrix completion based approaches to drug-target interaction prediction has focused on the integration of two sets of spaces, genomic and chemical, and mapping the two categories of information into a joint space for interaction prediction [36]. Later, this work was expanded by taking into account pharmacological insert information [37]. Other approaches developed for this problem with successful results included supervised learning [1], Gaussian interaction profile kernel approaches [31], and twin-kernel (genomic and chemical) Bayesian matrix factorization [8, 9]. Probabilistic matrix factorization has also been successfully applied to the drug-target interaction problem [2, 3]. Kshirsagar *et al.* have used the low rank nature of biological interaction matrices to identify host-pathogen interactions [17]. Motivated by the success of these algorithmic efforts, our approach relies on efficient latent factor modeling of a large tensor.

Thus, we present here a novel approach that aims to harness these advances in machine learning systems to perform sample-efficient testing of novel drug-disease interactions, using transcriptomic data to identify gene-cancer associations. To achieve this goal, we first build a tensor representing drug-target-disease associations, and then predict new associations through latent variable decomposition. Our model returns a list of experiments sorted by the algorithm’s confidence of their success, thus dramatically reducing the number of experiments leading to the first hit. Since the hit discovery efficiency is dramatically improved in this manner, the algorithmic ideation approach can potentially be used to run a *de novo* drug discovery pipeline on human tumor samples. We hope that the work described here contributes to creating an impetus for algorithmic and rational drug discovery pipelines instead of those that rely on brute-force screening.

Additionally, we would like to recognize the benefit of open source software in realizing the technical aspects of the approach we described here. We have used the Python/NumPy/SciPy stack, Numba from Continuum Analytics, the scikit-tensor library, the ChemFP Python library, and the Pandas data analysis library in our implementation. In recognition of the importance of open source code we release our code base under the GNU Lesser General Public License v3.0, both to allow the community to access our code as well as to spur other researchers in the community to develop their algorithmic approaches on this critical problem.

## 2 Methods

Our method entails the construction of a drug-target-disease associations tensor, and novel association prediction in that tensor. We first extract meaningful drug-target-disease tuples from the public TCGA and STITCH databases, which we use to seed a three dimensional tensor. We then perform PARAFAC tensor factorization on this tensor to infer relationships between drugs and diseases. These inferred relationships consist of a drug-disease pair, and a set of gene targets by which data indicates they potentially interact. We then evaluate these predictions by querying on held-out public data. The results of the evaluated predictions are then used to update the training data for the model, increasing future prediction accuracy. We have also implemented a passive version of the method to demonstrate the benefits of active learning, in line with previous work on active learning in drug discovery [33].

**Table 1:**
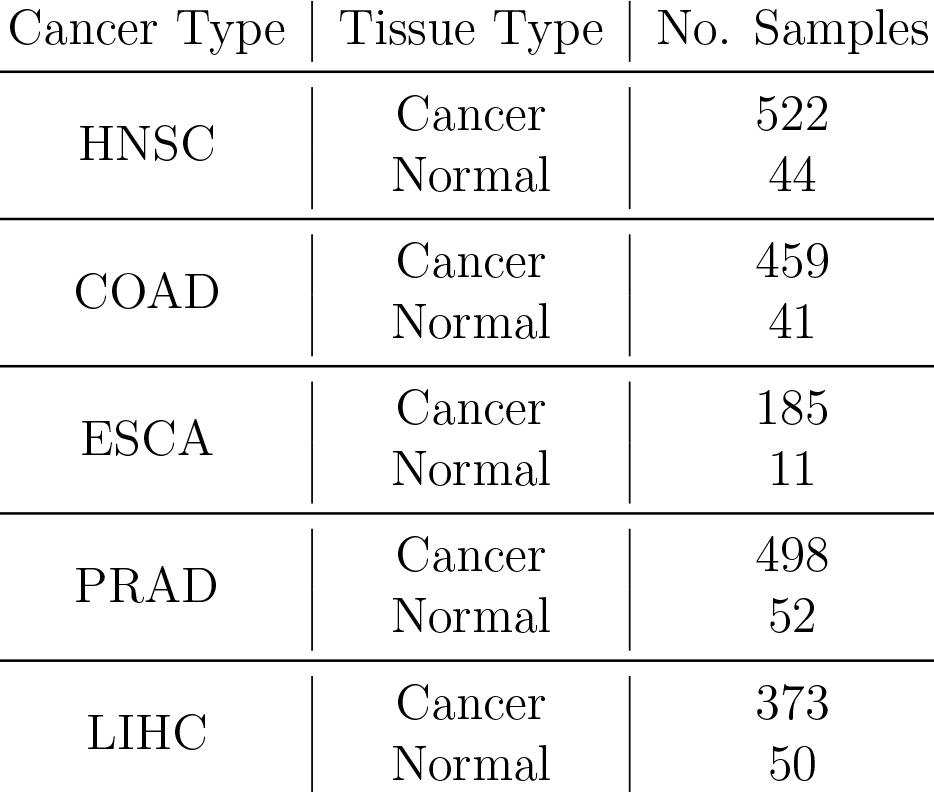
The number of TCGA samples across cancer and tissue types.

### 2.1 Generating drug-target-disease triplets from public data

To develop a training and testing set for evaluating our algorithm, we harness existing public data. We first identify a set of disease-target pairs using TCGA (The Cancer Genome Atlas) [34] data. We then cross-reference this information with the drug-target pairs in the STITCH database [19].

#### 2.1.1 Generating disease-target pairs from TCGA

We developed a measure of gene differential expression using transcriptomic profiling data (RNASeq v2) from five different TCGA cancer types from both cancer and normal samples, with the cancer types and the number of samples shown in Table 1. For each cancer type, we perform a t-test comparing the normal and cancer sample readings for each gene. We have manually selected a highly stringent p-value cutoff separately for each cancer. The cutoffs are highly stringent: even the least stringent threshold is lower than a Bonferroni-corrected threshold of 0.001 (which corresponds to a p-value cutoff of 5e-8). Cutoffs were selected to maintain a roughly equal number of genes selected per cancer. The distribution of the significance values and the selected thresholds are shown separately for each cancer in Supplementary Figures 1-5. Each significantly perturbed gene is assigned a score equal to the negative log of the p-value of their perturbation (and thus higher scores imply more confidence in their perturbation).

#### 2.1.2 Generating drug-target-disease triplets

We then cross-reference our disease-gene links with a set of drug-gene links derived from the STITCH database. We use all STITCH interactions between chemicals and human genes with reported confidence above 40%. This score from STITCH is a combination of experimental evidence, database curation, literature text mining, and predictions based on chemical structure [19].

To join the TCGA and STITCH data, we first represent it as a tripartite graph as shown in Figure 1a. This graph contains nodes for diseases (i.e., cancer subtypes), drugs, and genes. Edges connect drugs to genes (using STITCH data), and diseases to genes (using TCGA data). Each edge weight corresponds to the score for that interaction from the given database. For each pair of drug *D_i_* and disease *I_j_*, we calculate a score:

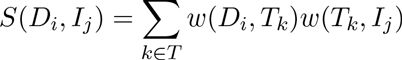

where *w*(*D_i_*,*T_k_*) and *w*(*T_k_*, *I_j_*) are the edge weights from *D_i_* to *T_k_* and *T_k_* to *I_j_* respectively, for each gene target *T_k_*. We then select a threshold score *S* such that all drug-disease pairs with a score above *S* are chosen as putative interactions. For the results reported here, we have selected *S* to be such that the most confident 10% of the predicted drug-disease interactions remain in *S*. (Our algorithm is robust to the selection of this cutoff threshold, see Supplementary Figure 7).

For each of these drug-disease interactions that we select as our positive set, we furthermore know the set of gene targets through which both the drug and disease show activity (all targets *T_k_* with weights *w*(*D_i_*, *T_k_*) and *w*(*T_k_*, *I_j_*) greater than 0). We consider these drug-target-disease triplets to represent specific genes through which the selected drug might modulate the selected disease.

### 2.2 Learning on a drug-target-disease tensor

Now that we have developed a set of canonical drug-target-disease interactions, we can perform learning to discover new putative drug-target-disease interactions. Discovering these novel triplets may result in possible drug hits targeted against specific cancer subtypes, while simultaneously providing the gene target(s) through which they interact.

**Figure 1:**
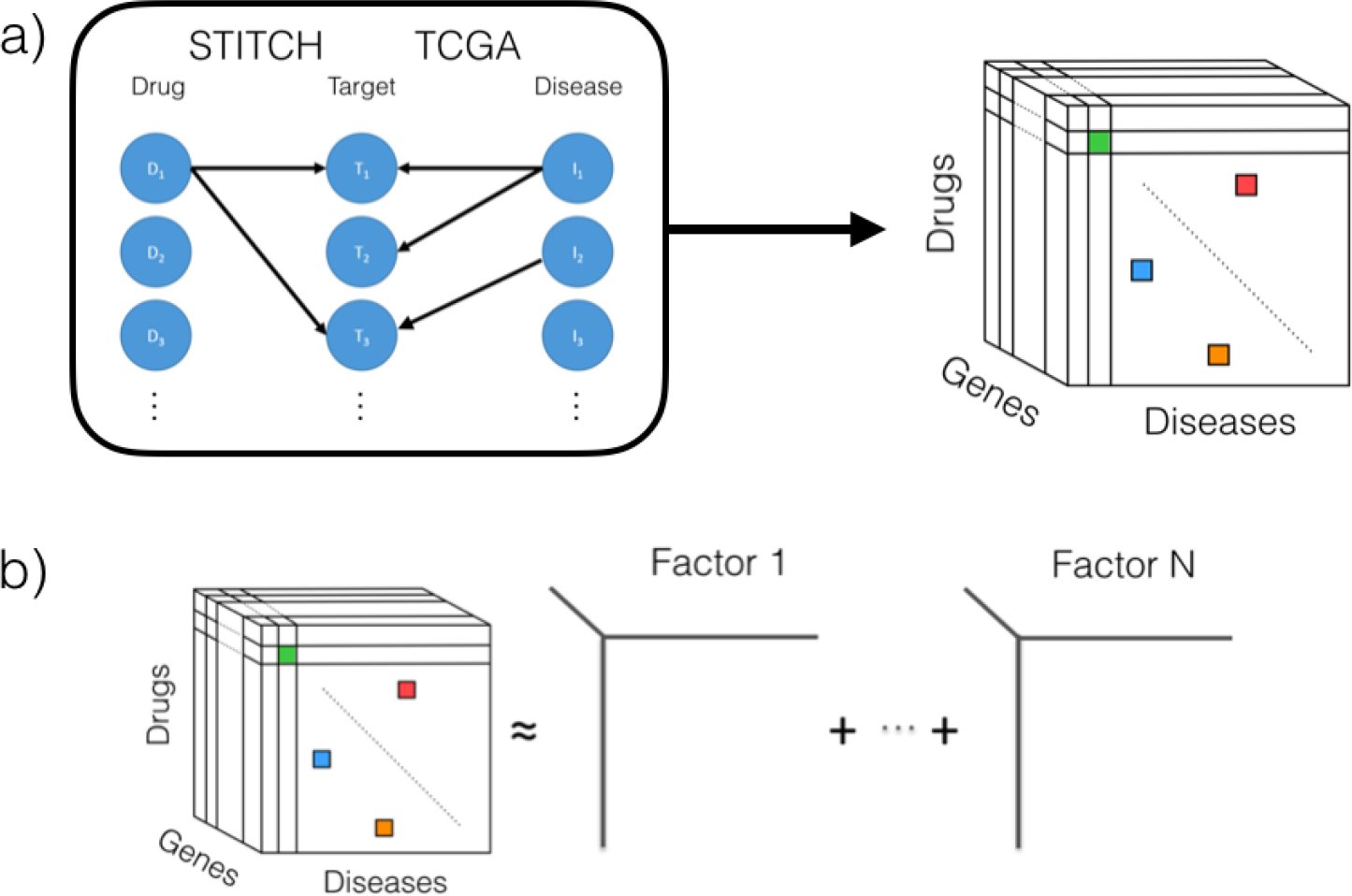
Overview of the method. (a) We identify drug-target-disease connections using STITCH for connecting drug candidates to targets, and TCGA for connecting targets to diseases, and construct a tensor by cross-referencing the data. (b) We use PARAFAC decomposition to create a latent factor model that reconstructs the tensor as a summation of the outer product of factor vectors, thus enabling the completion of the tensor from sparsely known entries.

We represent our data as a three dimensional tensor, with the dimensions being drug, target and disease. We take each known triplet and fill in their cell of the tensor with 1. Unknown triplets remain at 0; hence the value of each cell is based on the confidence of it being a true interaction.

Once all known entries have been input into the tensor, we perform tensor completion. Having evaluated several approaches, we use the venerable PARAFAC approach [11] due to its relatively rapid runtime and appropriately sized latent factor space. In PARAFAC, the tensor is decomposed into a number of latent factors as shown in Figure 1b. A parameter *N* is chosen, representing the size of the latent factor space.

We initialize the latent factors using eigen-decomposition of the unfolded tensor, which provides a good and deterministic starting point. These latent vectors can also be initialized randomly, and using an ensemble of such initialized models would also likely work well (however it would come at significant computational cost, as the model training would have to be repeated for every member of the ensemble at every update step of the active learner).

After initialization, we use alternative least squares to optimize the fit of the model to our data. We repeat the alternating process until the model converges (i.e. the fit of the model changes by a very small amount, 1e-4). Once learned, we can use the model to predict all entries of the tensor as follows:

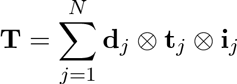

where **d***_j_*, **t***_j_*, and **i***_j_* respectively denote the drug, target, and disease factor vectors corresponding to the *j*-indexed factor of the decomposition model, and ⊗ represents the outer product. Thus, given a drug dimension of size *I*, target dimension of size *J* and disease target dimension of size *K*, PARAFAC learns (*I* + *J* + *K*) ∗ *N* parameters.

### 2.3 Performing learning on drugs with no known interactions

To be able to conduct *de novo* cancer drug discovery, it is important for the algorithmic approach to be able to generalize to drugs with no previously known interactions. However the standard PARAFAC algorithm cannot make any inferences on columns/rows with no known entries. For these drugs without any known interactions, we use their chemical structure to identify drugs with known interactions that have ¿95% chemical similarity (as calculated by ChemFP v1.1 OpenBabel FP4 fingerprints). We carry over the interactions of these highly similar drugs, but only after ‘dampening’ them by multiplying with a small coefficient 0 < *C* < 1, as follows:

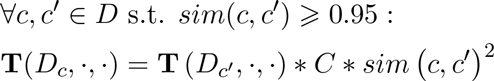

where *sim*(*c*, *c*′) is the chemical similarity between unknown (entry-less) drug *c* and known drug *c*′, **T** is the tensor, and *D* is the set of all drugs. We set the value *C* to 0.25 in our implementation. The motivation for using a dampening coefficient, *C*, is to reflect the intuition that chemical similarity is an imperfect indicator of functional similarity for drugs since drugs with high chemicals similarity can end up having different interaction dynamics. For example, an extra methyl group can render a drug incapable of fitting to the same binding pocket as one without the methyl group, yet these two structures would likely have very high similarity. It further allows us to reduce the weight of these imputed interactions compared to observed interactions from the public data. We multiply *C* with the square of the similarity to expand the difference between the similarity scores. Since chemical similarity above 95% is required for out cross-drug inference, this coefficient has a range of [0.9025, 1]. Where a compound has multiple analogues we carry over the interactions of all of its analogues, and if multiple analogues overlap on the same target-disease, then we arbitrarily select the last one as a tie-breaker - since only the similarity varies among ties, and that variation is restricted to such a small range, tied scores are always very close.

### 2.4 Software

The software used in this work is implemented in Python and available in an open source git repository at https://bitbucket.org/aiinc/drugx. It is licensed under the GNU Lesser General Public License v3.0.

## 3 Results

We have described here a novel algorithmic pipeline that allows us to predict likely interactions between drugs and diseases, as well as the gene targets through which they interact. We demonstrate the ability of algorithmic ideation to drive *de novo* interaction discoveries through four facets:

i. Our approach can predict hits efficiently
ii. The low-rank assumption intrinsic in a factorization model holds
iii. Our approach can work effectively in a low data (sparse) regime
iv. Our approach can infer interactions on novel chemicals

The results section is organized into subsections that discuss our results for each of these claims.

### 3.1 Drug-target-disease association prediction performance

We evaluated the algorithm using a temporal cutoff of gene-compound interactions: for training, we use the compound interaction data in STITCH 3.0 [18]; for testing, we use the new compound interactions found in STITCH 4.1 [19]. Effectively, this amounts to a temporal cutoff where the training data was collected before 2012, while the test data was collected between 2012 and 2015. Given this data, we constructed a 18.6-billion entry space, consisting of 5 diseases, 292,631 compounds, and 12,737 genes with 37,800 entries in the training set and 39,805 entries in the test set. Transcriptomic information was considered always available.

The results of our algorithm on this test can be seen in Figure 2. We define a ‘hit’ as any disease-drug-gene tuple that we propose for experimentation that is present in our test set. The predictions are made in 10 batches, with each batch comprised of 5,000 predictions for a total of 50,000 predictions. Our algorithm is highly insensitive to the choice of these batch number and size parameters as shown in Supplementary Figure 8. In the evaluation of our method, we follow previous work on active learning in drug discovery [33, 2] and present the results from both the active and passive versions of our method. Therefore we compare our active learning approach to a passive version of the same algorithm, as well as high throughput screening (random). The active versus passive comparison demonstrates that the tensor factorization based approach to the problem of predicting targeted antineoplastics is the main driver of hit discovery efficiency. We find that our algorithmic approach cumulatively performs 48,204-fold better than HTS. The batch with the best performance achieves 124,351-fold improvement and the batch with the worst performance still achieves 15,543-fold improvement. This demonstrates that the presented algorithmic ideation always outperforms standard screening approaches 10^4^-fold.

**Figure 2:**
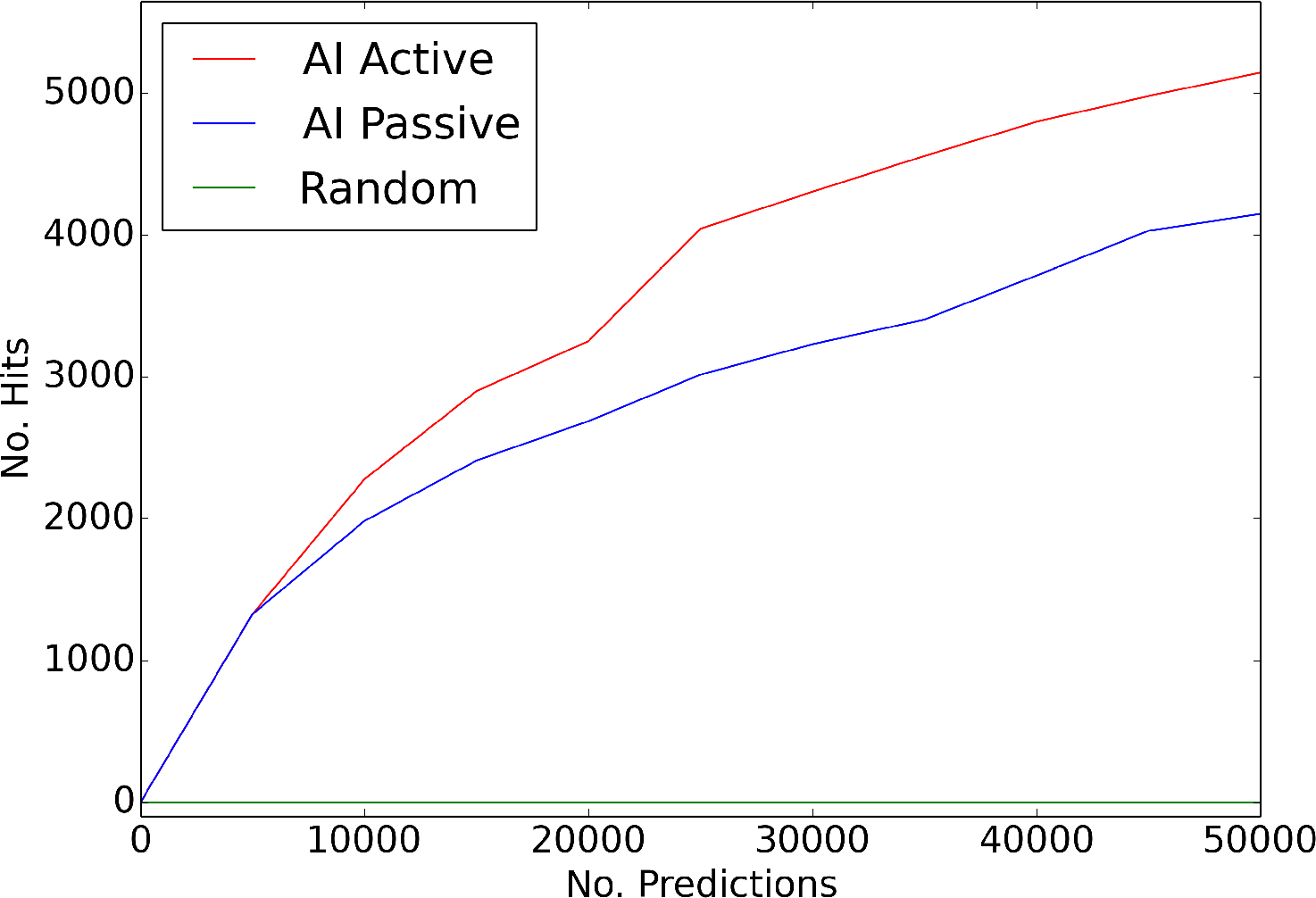
Predictive performance of the presented algorithmic ideation (AI) approach with active learning (red) and passive learning (blue). Brute force screening (random; indicated in green) is used to model the hit discovery performance of high-throughput screening. Training and testing were performed with a temporal cutoff using two versions of the STITCH database: STITCH v3 (training, data collected up to 2012) and STITCH v4.1 (test, data collected from 2012 to summer 2015). The initialization and the optimization are also deterministic, hence the results are deterministic. Algorithmic ideation significantly outperforms the brute force method in terms of hit discovery efficiency. Predictions done in 10 batches of 5000 predictions.

### 3.2 Testing the low rank assumption

The use of latent factor decomposition inherently assumes that the space is low rank, therefore it is important to make an effort to check that this assumption is valid. It is impossible to directly validate or refute the low rank hypothesis on this specific tensor, as calculating the actual rank of the tensor is impossible without knowing the full tensor, and it is unfeasible to complete the entire tensor (in fact that is precisely the reason we want to algorithmically complete this tensor). Hence we have estimated the inherent rank of the tensor through testing the method’s performance at different rank values with the idea that if the space is low rank then the performance should also peak at low rank: if decomposition rank is too low, this would lead to low performance due to not being sufficiently descriptive, whereas too high a rank would lead to overfitting and thus degrade performance. The results of this test are shown in Figure 3. Too low rank values cripple the performance, in fact when the rank is 1, the method effectively discovers hits only on the first batch and then becomes exhausted (Supplementary Figure 6). Slightly higher but still very low rank values, such as 7 to 70, lead to high performance with a plateau between 15 and 60. Higher rank values, such as 120, lead to a loss of performance due to overfitting. Peak performance occurs at rank 10. We have compared active and passive implementations of the method and the 10^4^-fold improvement persists in either scenario, yet active learning consistently outperforms passive (as to be expected). This demonstrates that the key factor behind the dramatic efficiency improvement is the algorithmic approach.

**Figure 3:**
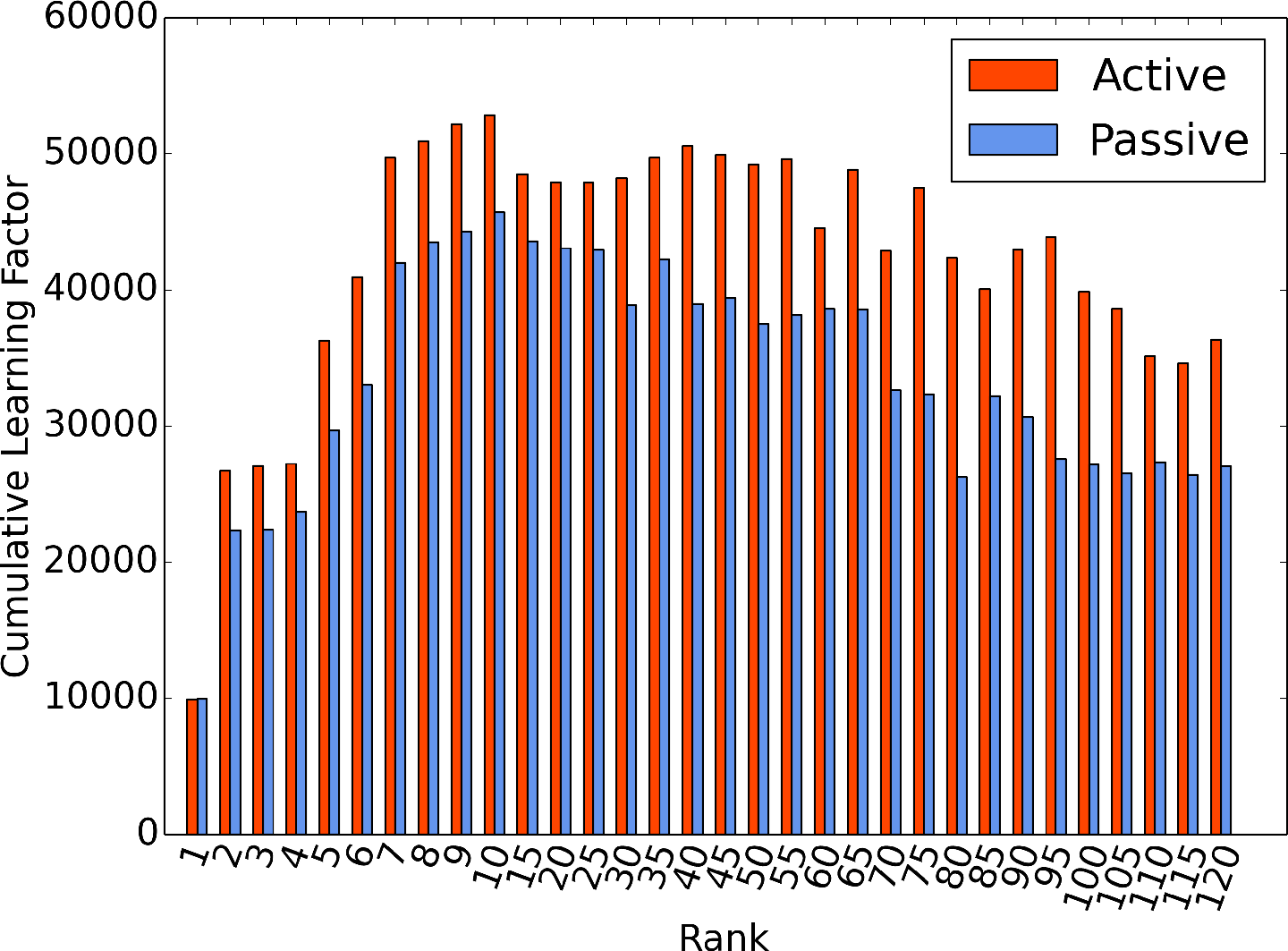
Predictive performance as a function of latent factor decomposition rank, with active (red) and passive (blue) implementations of the presented method. Results suggest that the tensor the low rank assumption is reasonable: at very low rank the method performs very poorly, then the performance peaks, and yet higher rank values lead to loss of performance. Active and passive approaches perform similarly, with active learning consistently demonstrating a small yet significant advantage.

At this point we would like to underscore that we did not conduct the rank scan as a hyperparameter search, but solely for the purpose of testing the low rank assumption. We did not pick rank 10 (where performance peaks) for use in all the other results in this paper, because that would constitute train-test contamination as we would effectively be using the testing set in the identification of hyperparameters of the method [5]. Instead we could either implement a cross-validation routine that splits the training data into training and validation sets and picks the top performing rank parameter separately every time the learner is called; or we could simply *ab initio* select a value that we think is reasonable. We have opted for the latter, mainly to demonstrate that the method and the conclusions presented here (namely that this type of algorithmic ideation confers 10^4^-fold hit discovery efficiency) are highly robust and do not require hyperparameter fine-tuning.

An alternative way to think about the representation of a 10dimensional latent factor decomposition is that this representation models a space of (*I* ∗ *J* ∗ *K*) with (*I* + *J* + *K*) ∗ *N* parameters where *I*, *J*, *K* are the sizes of the dimensions, and *N* is the number of latent factors. Thus we can write the number of tensor entries modeled by every single parameter as 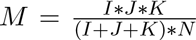 and if we plug in the values for our tensor we can calculate that *M* = 6103, which means that each parameter models about 6,103 entries of the tensor. Thus the 10-dimensional representation is low-rank in the sense that it is efficient in describing the tensor with few entries, however it can also achieve sufficient descriptive power to allow us to effectively discover hits at 10^4^-fold effectiveness compared to brute-force screening.

This sparse representation is due to most members of the same class being highly similar to each other. For example almost every drug class has multiple members (SSRIs, statins, etc.) and even though each drug is represented by a separate matrix, they are highly similar. Likewise cancers share aberrations in similar pathways - every cancer type does not create an entirely unique growth aberration. Targets are the potential exception, since each drug target protein can have unique characteristics but even then one can expect genes that are regulated with the same programs to change with similar trends. Therefore knowledge about one member of the class (drug, target, or disease) is often transferable to other members. Hence we think that the decomposition would be low rank even if significantly more data were to be added about other drugs or cancer types.

### 3.3 Predictive performance in low data regime

Beyond simply demonstrating that our approach finds more hits than HTS, we have also investigated several other properties of our algorithm that demonstrates its suitability to realistic, information-poor testing scenarios. This is important because drug-target-disease associations are difficult to compile, and will never be available in a large quantity compared to the size of the space of possible interactions. Thus, the approach should demonstrate utility even when only highly sparse training data is available. In Figure 4, we show that our approach is robust to scenarios with minimal training data by holding out a randomly selected fraction (up to 90%) of the training data. To evaluate the effect of removing different subsets of data, the tests were repeated 8 times in each condition. Figure 4 shows that the results are mostly robust across these repetitions. The scenario with 100% of the training data is equivalent to the setup discussed in Section 3.1, with both the training/testing data and the method deterministic, hence variance zero. To give a sense of the scale of the training data compared to the total space, in the most extreme case tested, we used only 10% of our training data, which results in fewer than 4,000 known interactions being used. Even in this extreme scenario, we observe close to 10^4^-fold improvement. As the training data used increases, so does the performance which is exactly as expected. Active learning consistently confers an advantage, however the passive implementation also benefits from the additional boost in data which shows that the ability of the presented approach to effectively utilize the available data is the key driver of performance, with the active component offering a secondary advantage.

**Figure 4:**
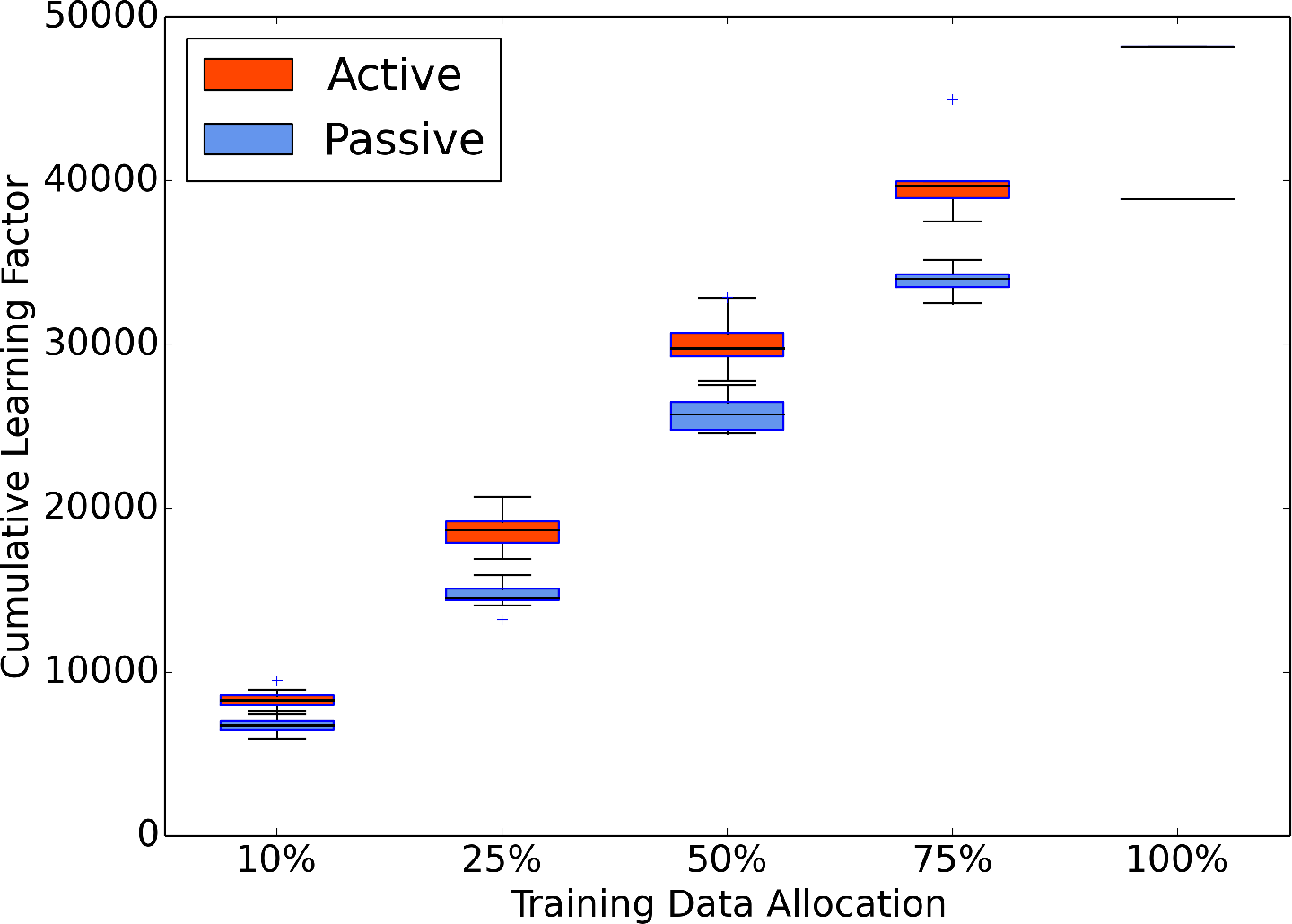
The learning factor (fold-improvement over random) as a function of training data constriction for active (red) and passive (blue) implementation of the proposed method. At *x*% training data allocation, 100 — *x*% of the data in the training set is randomly removed. Since data is randomly removed, the train/test sets are stochastic and the results can vary, hence at each point the experiment is repeated 8 times and the distribution of the maximum learning factors are reported as a boxplot: center line indicates median, top and bottom lines of the rectangle represent the first and third quartiles respectively, whiskers extend to any points within 1.5 standard deviations, and outliers are indicated with a ‘plus’ sign. 100% training data is equivalent to using all the STITCH v3 data and is thus deterministic (there is zero variance in the results). The algorithmic ideation approach effectively performs even in low data regimes.

**Figure 5:**
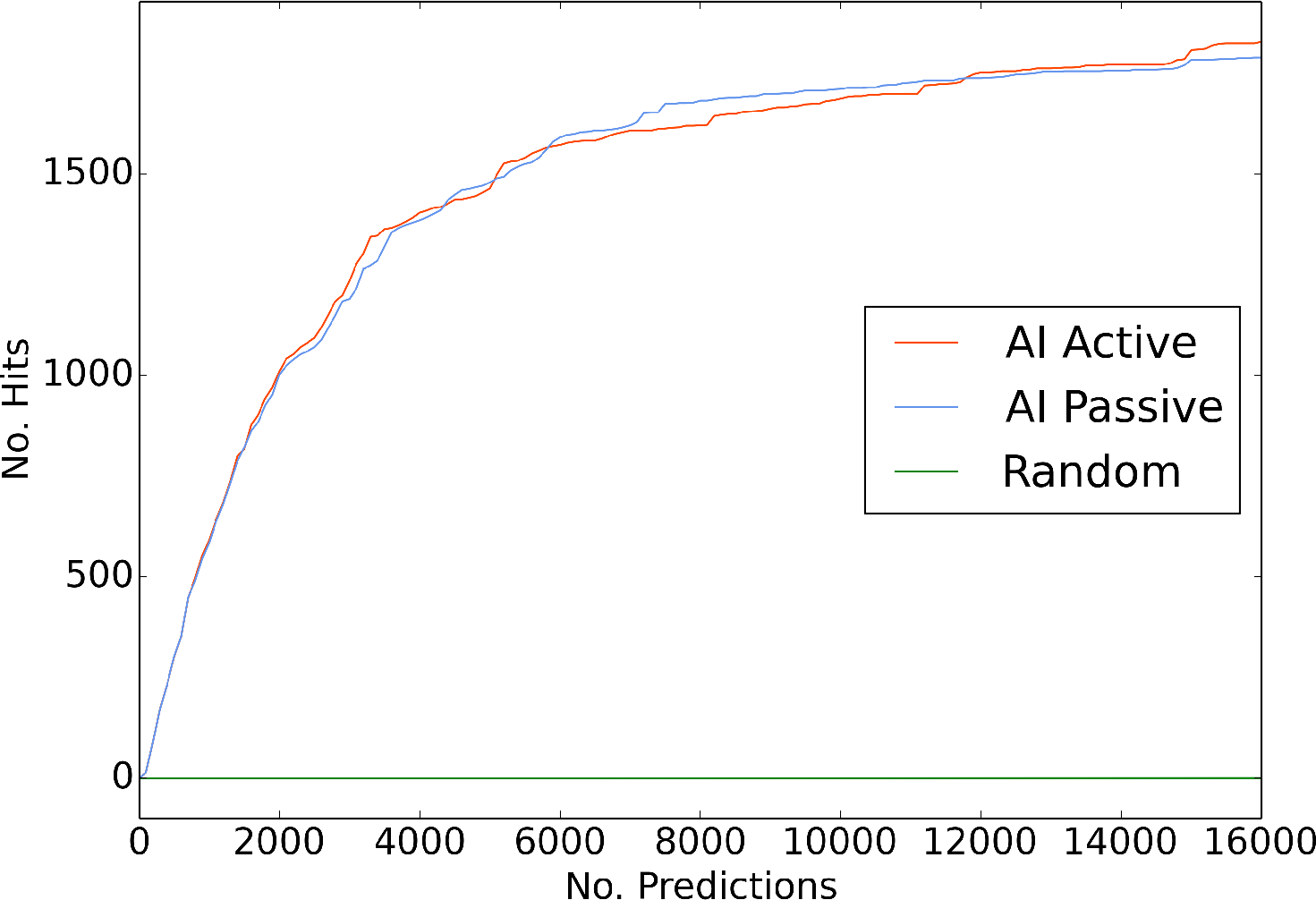
Predictive performance on *de novo* drugs. The x-axis shows the number of predictions made, and the y-axis shows the number of hits discovered in those predictions. The results from both the active (red) and passive (blue) implementations are shown. The green line shows brute-force arbitrary order compound selection, (labeled random). Even in the more challenging scenario with no known interactions for the test set chemicals, the method still outperforms brute-force testing, albeit at a rate of 10^3^-fold instead of the rate of 10^4^-fold observed when some associations are known about the drugs.

### 3.4 Predictive performance on *de novo* drugs

To perform *de novo* discovery, it is important to be able to make predictions on chemicals for which no interaction data is known. Our method is capable of reasoning in this scenario, as we use the distance metric described in Section 2.3 to infer interaction information for compounds with only chemical structure information. To test the *de novo* scenario, we separate the training and test data using a cutoff similar to that in Section 3.1, with the difference that we separate the data so that we train on *all* of the known interactions of compounds that had any interactions known in STITCH 3, while we test on compounds for which interaction data is only found in STITCH 4. Hence, for all compounds in our test set, the training set has no entries in the corresponding plane of the tensor.

In Figure 5, we show the number of hits resulting from the predictions of our method in 160 batches of 100 predictions. The results of 50,000 predictions in various batch size and batch number combinations are available in Supplementary Figure 8, and they show that the algorithm is highly robust and performs similarly with different choices of batch size and number. Even at the first batch we are able to identify hits at a 4,165-fold improvement over HTS. The cumulative learning rate (fold improvement over brute force) ranges between 1,233 and 19,040 with a median of 2,262. Since predicting in the *de novo* chemical space is a much harder problem than predicting on chemicals for which some interaction data is already known, the fold improvement falls from 10^4^ to 10^3^. However the key point we wish to emphasize here is that the algorithmic ideation approach still significantly outperforms the established brute force testing of chemicals, and is capable of effective learning even on *de novo* chemicals. These results show that our algorithmic ideation can use the public data for training and generalize to any chemical space. Therefore the approach proposed here can be utilized to run a screening on any chemical library.

## 4 Discussion

Currently there exist two different types of assays to test anticancer drug effectiveness. On the one hand there exist functional assays test few drugs but on human tumor tissue/cells thus have very high relevance to human disease [7], almost exclusively used for diagnostic purposes, specifically to identify the best drug among the currently available treatment options for a given patient. On the other hand, *de novo* drug discovery requires high-throughput screening over millions of chemicals and even the most relevant of these assays are conducted on monotypic cell lines in test tubes - missing critical aspects of relevance to human disease such as heterogeneity and microenvironment [25].

We have demonstrated here a novel technique to harness public data to create an active learning-based ‘algorithmic ideation’ system that proposes linkages between drugs, diseases, and the genes through which they interact. By increasingly the efficiency of hit discovery in screening by a factor of 10^4^, we believe that our approach renders it possible to use functional assays for *de novo* drug discovery for the first time.

It is worth mentioning that the method performance seems to plateau after discovering a certain number of hits. This shows that with the latent factor decomposition based approach, there are only so many unknown novel associations that can be predicted. However the critical point to note here is that the method’s predictions are not tested in actual experiments - when the method asks for the testing of an association, if that drug-target interaction has not been reported in STITCH, we consider that a miss. The interactions reported in STITCH are likely to be true (otherwise they would not be reported) however the converse is not true: there are likely to be many more interactions between the chemicals and targets in our testing space that have simply not been discovered. Therefore, our results represent a lower bound on the performance of our predictions - the actual performance cannot be lower, but is likely to be higher. We argue that the exhaustion of hits after a few thousand hits is due to the fact that because the model’s predictions are not actually tested, the model cannot infer the actual space and simply a smaller and less diverse subspace.

## Acknowledgements

We would like to acknowledge the support for our company from the Project Olympus of Carnegie Mellon University.

